# Scop3P: a comprehensive resource of human phosphosites within their full context

**DOI:** 10.1101/684985

**Authors:** Pathmanaban Ramasamy, Demet Turan, Natalia Tichshenko, Niels Hulstaert, Elien Vandermarliere, Wim Vranken, Lennart Martens

## Abstract

Protein phosphorylation is a key post-translational modification (PTM) in many biological processes and is associated to human diseases such as cancer and metabolic disorders. The accurate identification, annotation and functional analysis of phosphosites is therefore crucial to understand their various roles. Phosphosites (P-sites) are mainly analysed through phosphoproteomics, which has led to increasing amounts of publicly available phosphoproteomics data. Several resources have been built around the resulting phosphosite information, but these are usually restricted to protein sequence and basic site metadata. What is often missing from these resources, however, is context, including protein structure mapping, experimental provenance information, and biophysical predictions. We therefore developed Scop3P: a comprehensive database of human phosphosites within their full context. Scop3P integrates sequences (UniProtKB/Swiss-Prot), structures (PDB), and uniformly reprocessed phosphoproteomics data (PRIDE) to annotate all known human phosphosites. Furthermore, these sites are put into biophysical context by annotating each phosphoprotein with per-residue structural propensity, solvent accessibility, disordered probability, and early folding information. Scop3P, available at https://iomics.ugent.be/scop3p, presents a unique resource for visualization and analysis of phosphosites, and for understanding of phosphosite structure-function relationships.

## Introduction

Post-translational modifications (PTMs) are typically the result of the addition of a small molecule to one or more residues of a protein ^1,2^. PTMs can be reversible as well as irreversible, with more than 200 PTMs currently identified. Protein phosphorylation, a reversible PTM, is one of the best studied PTMs and is involved in many regulatory processes ^3,4^. Protein phosphorylation is regulated by three core machineries, namely kinases, phosphatases, and proteins which recognize the phosphorylation signals/P-sites ^5^. Kinases, one of the largest protein families, can be considered as the writers of protein phosphorylation: they add a highly negatively charged phosphate group to the side chain of serines, threonines, and tyrosines and less frequently to cysteines and histidines ^6^. These phosphorylation signals can be recognized by phospho binding proteins – the readers – which interact with phosphorylation signals and proteins^7^. Phosphatases, the erasers of the phosphate group, function opposite to kinases: they remove the phosphate group from phosphorylated residues ^7,8^.

Several studies have attempted to differentiate between functional and non-functional P-sites based on their evolutionary conservation^9,10^, kinase specificity ^11,12^, PTM cross-talk or based on their interactions ^13^. However, P-site conservation may not be particularly useful to determine functional importance of a P-site as only a small fraction (∼35%) of functional P-sites were reported to be conserved ^14,15^, while some functional P-sites have been identified in poorly conserved regions ^11,16^. Most of the conserved but non-functional P-sites have been accumulated due to the off-target effect of kinases ^11^ in disordered regions of proteins as these are more accessible to kinases than ordered regions are ^17^.

The dominant means of discovering novel P-sites is mass spectrometry (MS) based phosphoproteomics, which has been the primary driver for the expansion of the known P-sites. With the increasing public availability of data from such proteomics experiments ^18^, several databases related to one or more PTMs in one or more model organisms have been established. For instance, O-GLYCBASE ^19^ is an information repository of the glycosylation status of proteins, while the Human Protein Reference Database (HPRD) ^20^ contains information on experimentally annotated PTMs of human proteins, and dbPTM ^21^ contains experimentally verified as well as computationally annotated PTMs. Besides these generic PTM databases, PHOSIDA ^22^, Phospho.ELM ^23^, Phospho3D ^24^, PhophositePlus ^25^, and database of Phospho-sites in Animals and Fungi (dbPAF) ^26^ are all databases that focus on P-sites specifically and contain information about the sequence and/or structural features of experimentally determined P-sites. Many of these sites are also reported in UniProtKB/Swiss-Prot ^27^, which contains both functional and structural annotations of proteins, but lacks direct access to important structural properties related to P-sites. Most of the abovementioned repositories collect and integrate a large number of P-sites from different sources or MS experiments, but provide very little or no information on the functional role(s) of these P-sites. Moreover, the phospho-peptides measured in different phosphoproteome experiments are typically identified with different search engines and different false discovery rate (FDR) thresholds, which may artificially increase the heterogeneity of P-sites when these are integrated into a database which has no information on the significance of the reported P-sites ^28^. This issue can be mitigated by analyzing the entire data set as one, thus allowing the control of the global FDR threshold ^29^.

Because of the difficult and time-consuming process of experimental identification of P-sites, several computational P-site prediction algorithms have been developed ^30–33^. These predictors are typically trained on data obtained from the above described public resources or on the observation of kinase specificity and sequence features to predict if a particular site can be phosphorylated. However, these predictors provide very little to no information on whether the site is functional or not. They also often neglect the importance of conformational specificity of kinases^34^ and of structural dynamics upon phosphorylation/de-phosphorylation ^35^, which can lead to incorrect or non-confident P-site predictions.

Because the function of a protein and the corresponding signaling cascades are highly correlated with protein structure, and because phosphorylation status can result in protein structural rearrangements ^36,37^, it is important to know where a phosphorylation is located on the protein structure to understand its possible regulatory role. Visualizing P-sites mapped onto available protein structures can provide such insight by presenting researchers with an overview of the spread of P-sites over the three-dimensional structure of a protein.

Beyond the structural context itself, there is also the biophysical context of a residue. Indeed, adding or removing a negatively charged phosphate group alters a residue’s electrostatic potential ^38,39^. This alteration may serve as a recognition site for phospho binding proteins but can also trigger conformational transitions in the phosphoprotein. Phosphorylation can moreover modulate the binding specificity of phosphoprotein binding proteins by offering a wide range of recognition patterns based on conserved amino acid residues close to the P-sites ^40,41^. Moreover, studies have shown an association between phosphorylation/de-phosphorylation events and order-disorder transitions that are in turn coupled with binding regulation ^38,42–44^.

It is thus clear that a thorough analysis of the biological relevance and possible role of a given P-site needs to take place against the full context of that P-site, which consists of the P-site localization in the protein, its structural characteristics, its experimental provenance, and its biophysical properties.

Here we therefore present Scop3P, a database of human P-sites in their full context. To do so, Scop3P provides annotation for all known human P-sites based on protein sequence, 3D structure, and biophysical predictions. Moreover, Scop3P is unique in that it also provides a reliability measurement for each P-site based on the frequency with which that phosphorylation has been seen across different phosphoproteomics experiments. Importantly, these phosphoproteomics results have been obtained by a uniform, large scale re-analysis of phosphoproteomics data from the PRIDE database ^45^, and have been filtered by a global FDR to high reliability. In addition, Scop3P contains secondary structural propensity (helix, sheet, coils), solvent accessibility, and biophysical properties such as the probability of being a disordered region, backbone dynamics, and functional information related to phosphoproteins. Importantly, every phosphoprotein is also annotated with early folding predictions ^46^, which give an idea of residues or regions that are crucial in the folding dynamics of the protein. By providing information on early folding regions/residues, Scop3P adds further unique information on whether or not a phospho acceptor residue is crucial in forming local structural elements that influence the final fold of a protein.

## Methods

To create Scop3P we collected and integrated all available P-sites from different data sources (Fig 2A-C, Table 1). First, we retrieved all available human P-sites from UniProtKB/Swiss-Prot ^27^(release-2018_02) by parsing UniProtKB/Swiss-Prot MOD_RES records. For every P-Site the evidence annotation (experimentally determined, or by similarity) and the associated reference information were also obtained.

**Figure 1.**
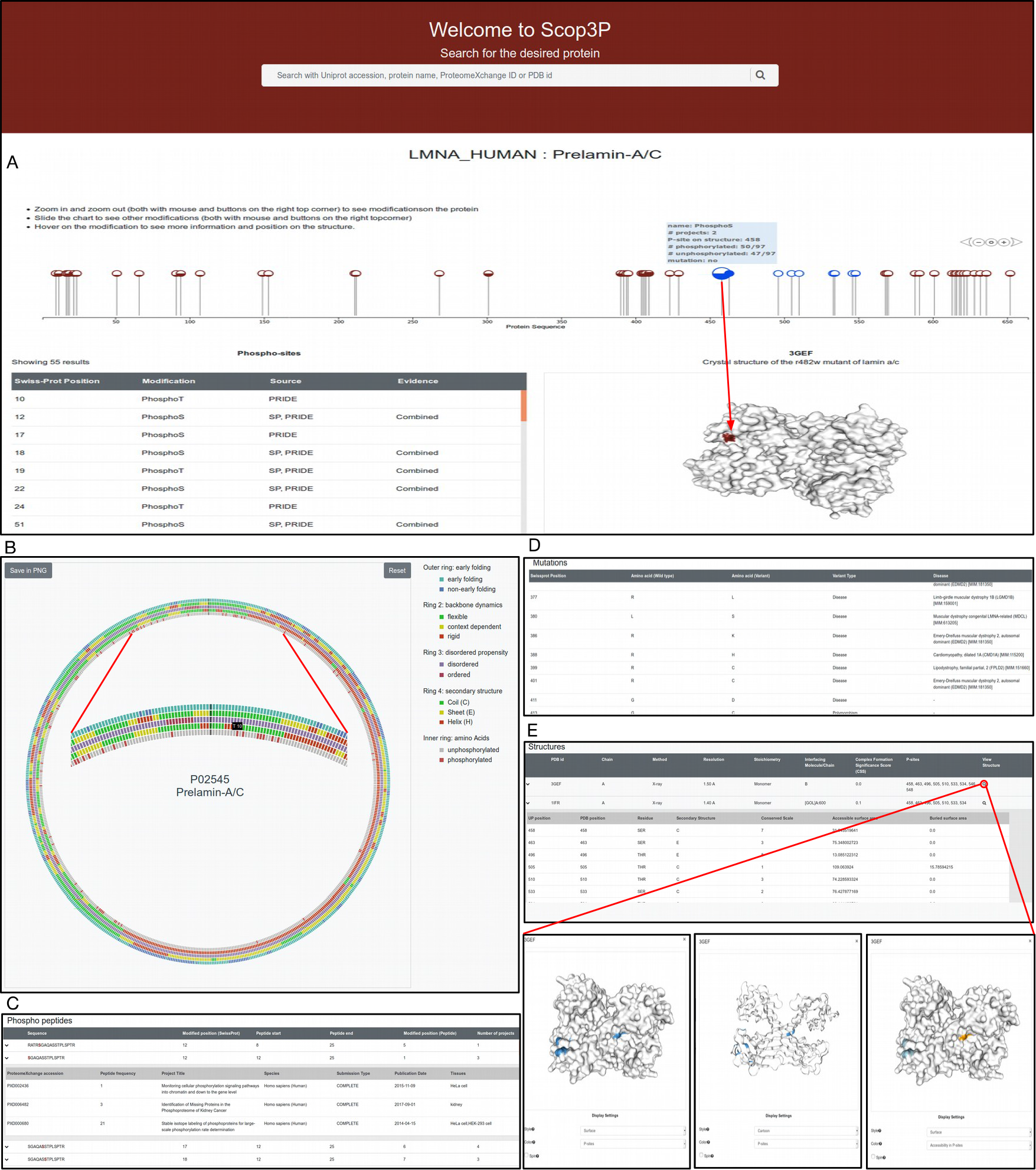
Scop3P web-interface reveals different context information at different levels for all phosphosites in a phosphoprotein. For a given phosphoprotein, Scop3P provides a variety of information in tables and interactive graphs. **A** Quick overview of all P-sites mapped onto the protein sequence as ball and stick. The color fill in the ball represents the number of different peptides identified for that particular site across PRIDE projects. This information is also presented as a table below. A tooltip over the ball will display additional information, while a quick preview panel on the right shows where that P-site is located on the protein structure. P-sites mapped onto the displayed structure (3GEF) are colored blue. **B** All predicted biophysical properties and secondary structures are presented in an interactive circular plot with different concentric rings. The graph can be zoomed to focus on a region of interest (inner panel). **C** Identified phospho peptides, source projects, and frequency information are presented in tabular form. Details about a project can be viewed by clicking the project ID (inner panel). **D** Mutation table provides information on amino acid variants, type of polymorphism, and associated diseases (if any). **E** Information related to structures, their secondary structure information, and P-sites mapped onto the particular structure are displayed in an overview table. Clicking the view structure icon will launch the structure viewer and shows P-sites mapped onto that particular structure. P-sites can be colored based on different properties.

**Figure 2.**
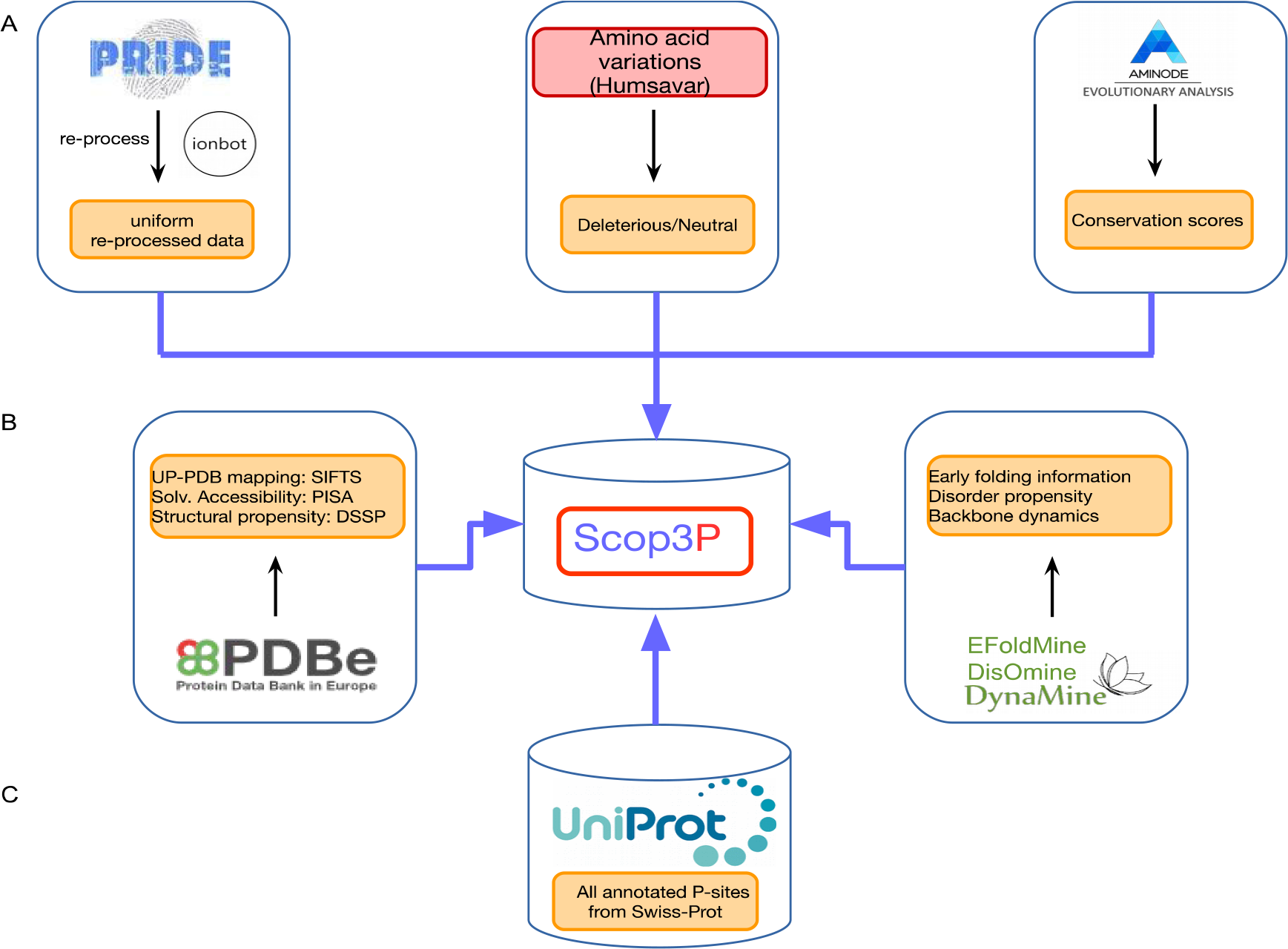
Scop3P data collection and integration flow. **A** Sequence level information like experimental phosphosites (P-sites) obtained from re-processed public proteomics data from PRIDE (37 projects), single amino acid variations and the associated disease and polymorphism details from Humsavar dataset and the amino acid conservation of P-sites from AMINODE were integrated onto the amino acid sequence of proteins. **B,C** All available P-sites from Swiss-Prot along with other structural information (secondary structures from DSSP, solvent accessibility from PISA) for all P-sites (Swiss-Prot+PRIDE) and predicted residue level biophysical properties of phosphoproteins (disorder and folding Propensities and backbone dynamics) were integrated on amino acid sequences. Sequence to structure mapping was done using SIFTS mapping.

**Table 1.**
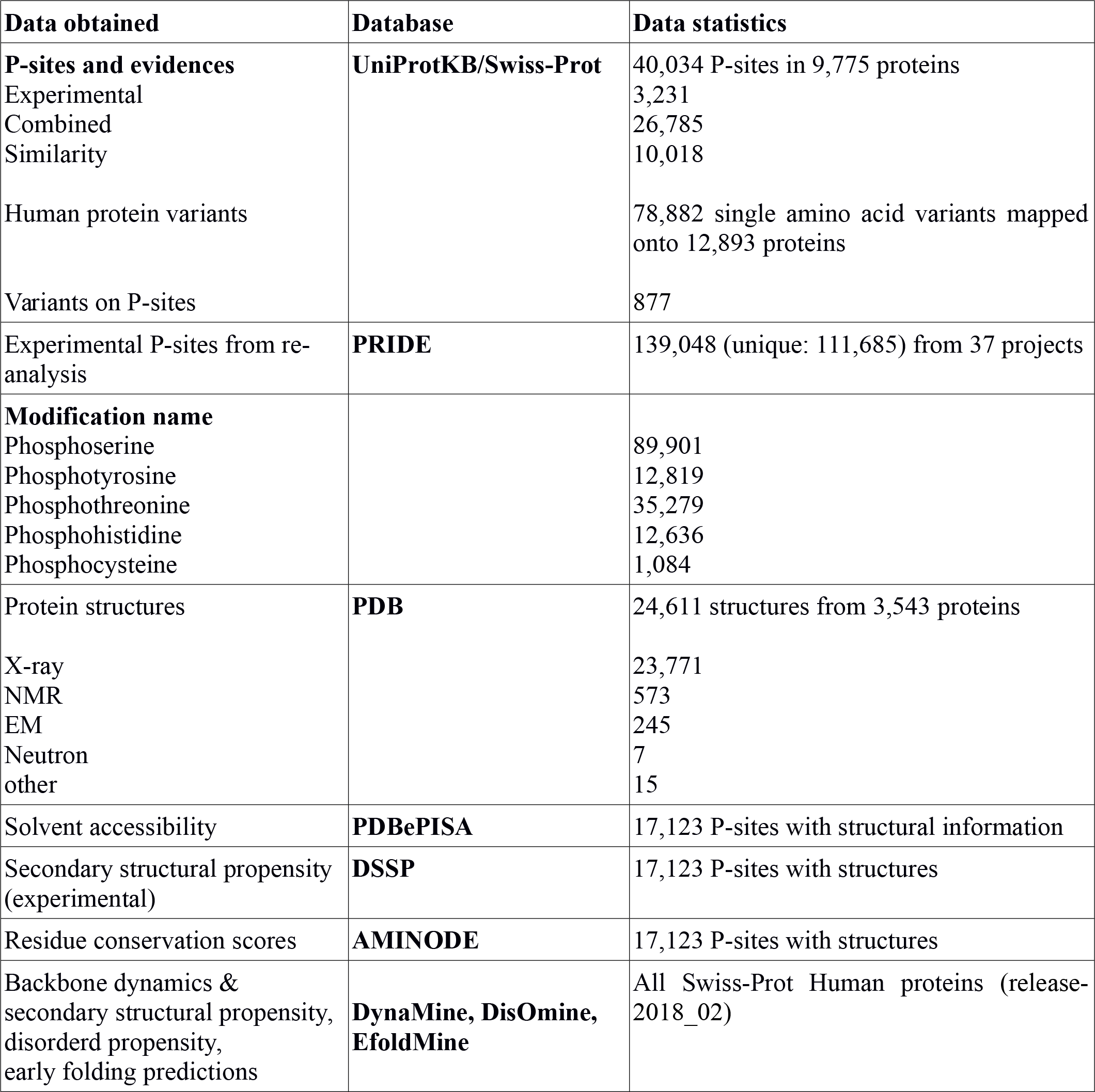
Data sources/tools and derived data integrated into Scop3P

| Data obtained | Database | Data statistics |
| --- | --- | --- |
| **P-sites and evidences**<br>Experimental<br>Combined<br>Similarity<br>Human protein variants<br>Variants on P-sites | **UniProtKB/Swiss-Prot** | 40,034 P-sites in 9,775 proteins<br>3,231<br>26,785<br>10,018<br>78,882 single amino acid variants mapped onto 12,893 proteins<br>877 |
| Experimental P-sites from re-analysis | **PRIDE** | 139,048 (unique: 111,685) from 37 projects |
| **Modification name**<br>Phosphoserine<br>Phosphotyrosine<br>Phosphothreonine<br>Phosphohistidine<br>Phosphocysteine |  | 89,901<br>12,819<br>35,279<br>12,636<br>1,084 |
| Protein structures<br>X-ray<br>NMR<br>EM<br>Neutron<br>other | **PDB** | 24,611 structures from 3,543 proteins<br>23,771<br>573<br>245<br>7<br>15 |
| Solvent accessibility | **PDBePISA** | 17,123 P-sites with structural information |
| Secondary structural propensity (experimental) | **DSSP** | 17,123 P-sites with structures |
| Residue conservation scores | **AMINODE** | 17,123 P-sites with structures |
| Backbone dynamics & secondary structural propensity, disorder propensity, early folding predictions | **DynaMine, DisOmine, EfoldMine** | All Swiss-Prot Human proteins (release-2018_02) |

### Re-processing of human phosphoproteomics data from PRIDE

We retrieved a list of all Human projects which are annotated to contain phosphorylations from PRIDE ^45^. Only those projects which are unlabeled and submitted as complete projects containing high resolution spectra files were considered (see appendix Table S1). In total 2032 RAW files containing 60.2 million (60,271,683) spectra from 37 different projects were retained for processing. These projects were typically originally processed with different search engines and different search settings. In order to obtain uniform data, we collected all ‘.RAW’ files from PRIDE and converted these to Mascot Generic Format (MGF) peak files using ThermoRawFileParser^48^ (Hulstaert et al, 2019). The resulting peak files were then searched against the human complement of UniProtKB/Swiss-Prot (release-2018_02, containing 20259 protein sequences) with the target/decoy approach using ionbot (https://ionbot.cloud/; ionbot is based on MS^2^PIP ^49–51^ and ReScore ^52,53^). Results were filtered at 1% FDR. The ionbot engine searches for all modifications listed in Unimod ^54^ on top of a set of user-defined fixed and variable modifications. The search settings were as follows: carbamidomethylation of cysteine was specified as fixed modification, and oxidation of methionine, phosphorylation of serine (S), phosphorylation of threonine (T), phosphorylation of tyrosine (Y), phosphorylation of cysteine (C) and phosphorylation of histidine (H) were set as variable modifications. Up to two missed cleavages per peptide were allowed. Only identified peptides with q-values <0.01 were considered for further analysis. In total, we identified 19.2 million (19,267,930) PSMs (peptide spectrum matches) with 1% FDR resulting in 151,719 Psites. The total search time was 12 days on a single Linux server with 24 cores and 30 GB of RAM memory.

### Structural properties of the phosphoproteins

For every human phosphoprotein for which at least one structure was available, the modeled segment of the UniProtKB/Swiss-Prot sequence in the protein structure was scanned to check if any P-sites were within range of that segment. If the modeled segment contained at least one such Psite, the corresponding PDB ^55,56^ structure was used to map and visualize all matching P-sites. Psites that fell in missing segments of structures were not considered for structural mapping. The oligomeric state of the protein structure, solvent accessibility of the P-sites, and exposure level (buried, exposed, or in an interface region) were obtained from the Protein Interfaces, Surfaces and Assemblies (PISA) server (also known as PDBePISA) ^57^. The interface details were obtained by taken into consideration the most probable quaternary structure as assigned by PISA. PISA predicts quaternary structures based on the interactions occurring in macromolecular crystals (pair of chain or ligand-chain interactions). The secondary structural assignments for P-sites with a matched structure were retrieved from DSSP ^58^. The eight-class classification of DSSP was grouped into three states (**helix (H):** ‘alpha helix, 3/10 helix, pi helix’, **strand (E)**: ‘extended strand, residue in isolated beta-bridge’, and **loop (C)**: ‘turn, bend and the rest’). Every structure is also annotated with its determination method, resolution, and stoichiometry details.

For all phosphoproteins, regardless of existing structure match, the three states of the secondary structural propensity (helix (H), coil (C), sheet (E)) were predicted using Fast Estimator of Secondary structures (FESS), which is a component of the FELLS method ^59^. Protein biophysical characteristics such as backbone dynamics, disordered propensity, and early folding properties were predicted using DynaMine ^60^, DisoMine (http://bio2byte.com/disomine), and EfoldMine ^46^, respectively. DynaMine predicts the residue-level backbone flexibility in the form of S^2^ values between 0 (highly dynamic) and 1 (stable conformation), which represents how restricted the movement of the atomic bond vector is with respect to the molecular reference frame. For DisoMine, the probability cutoff of 0.5 distinguishes the (predicted) ordered and disordered state of the protein. EfoldMine predicts the early folding (EF) propensity of amino acids based on local interactions, and as such provides insight into which amino acids are likely involved in early stages of protein folding and thus shape the folding landscape of that protein. The EF propensities and binary classification based on a 0.163 probability cutoff were used to distinguish between early folding and non-early folding residues.

### Conservation and variation of amino acids in phospho acceptor residues

Known amino acid variations for P-sites (phospho variants) or for sites in their close proximity may result in functional variants, e.g., through a change in kinase specificity, loss and gain of P-sites, and diseases ^61^. In order to map such variants on both sequence and structure, we retrieved all curated human missense variant details from the Humsavar dataset (release 12-Sep-2018) from UniprotKB/Swiss-Prot (Fig 2A, Table 1) that are classified as disease/polymorphisms/unclassified based on their role in disease. Humsavar data contains all manually curated single amino acid polymorphisms as retrieved from literature that are associated with diseases and phenotypes. In total it contains 72,960 variants of which 40% are associated with diseases. We also obtained the evolutionary conservation from AMINODE ^62^ for all P-sites mapped to a structure (Fig 2A, Table 1). These conservation values ranges from 0 (variable) to 1 (conserved).

### Database construction, integration and content

The web interface was developed using the Spring Boot framework. JQuery, Bootstrap and Tymeleaf were used as front-end technologies. Protein structures are visualized with the aid of NGL Viewer ^63^, and other protein visualizations such as the circular plot and ball and stick representation of P-sites on primary (one dimensional) amino acid sequences are developed through the D3 javascript library. Scop3P data is stored in a relational database running on MySQL 5.7.

Scop3P contains both sequence (Fig 2A,C) and structure (Fig 2B) information, for both phosphoproteins and for individual P-sites (Fig 1A-E). All obtained parameters were mapped to the amino acid sequences of the human proteins retrieved from UniProtKB/Swiss-Prot. Sequence to structural position mapping was done with the aid of SIFTS ^64^. P-sites which fell in the missing segments of available structures were not considered for structure mapping. Every instance with structure was annotated with secondary structural propensity and evolutionary conservation details as described earlier. Additional residue level biophysical properties such as DynaMine, DisoMine and EfoldMine predictions were annotated to UniProtKB/Swiss-Prot protein sequences. Moreover, to show the reliability of the P-sites, every P-site is annotated with the frequency of phosphorylation as found in the different phosphoproteomics experiments. As a second level of annotation, every Psite is annotated with the number of distinct peptides identified for that particular protein from different PRIDE projects, and every such peptide is then annotated with its ProteomeXchange ID, and its frequency across the different PRIDE projects.

In Scop3P, we aim to map all P-sites of a particular protein to all available three-dimensional structures. Thus, if a protein has more than one structure, all structures that contain at least one Psite are retained for mapping and visualization. For each P-site with 3D structure, the assembly and interface details such as the macromolecule chain where the P-site is present (main chain), accessible surface area (ASA), buried surface area (BSA), and information about crystal contacts like complex significance score (CSS) and interacting chains/ligands are also given.

### CSS score

PISA assigns a value from 0 to 1 for every complex (CSS) in the biological assembly. This value is calculated as a fractional contribution of the particular interface to the crystal assembly. Hence, in Scop3P, if a particular P-site is present in a multimeric protein (with chain XYZ) at chain X, only the interfaces with higher CSS value for the chain X are considered. For example: if the CSS value for interface XY is higher than XZ then this means that this complex – composed of chain X and chain Y – is the most probable biological assembly as predicted by PISA. Sometimes the interacting molecule will be a ligand which means that the ligand is fixed with the polymer during PISA prediction and the CSS for the main chain of P-site and this ligand is higher.

## Results

After populating Scop3P with all known phosphosites as obtained from UniProtKB/Swiss-Prot and the reprocessing of complete human phosphoproteomics experiments in PRIDE, the Scop3P database contains 15,728 phosphoproteins, covering 78% of the 20,259 human proteins in UniProtKB/Swiss-Prot. Together, these proteins contain a combined 151,719 P-sites (Table 1) of which 124,356 are unique P-sites (12,671 are unique to Swiss-Prot, 111,685 are unique to PRIDE) and 27,363 sites are shared. Total of 40,034 P-sites (experimental: 30,016, by similarity 10,018) are obtained from UniProtKB/Swiss-Prot annotations, and 139,048 by re-processing experimental data in PRIDE (Table 1). The distribution of all P-sites in Scop3P shows that 59.25% of P-sites are phosphoserine, 23.25% are phosphothreonine, 8.45% are phosphotyrosine, 8.32% are phosphohistidine, and 0.71% are phosphocysteine (Table 1).

The structural data in Scop3P contains 17,123 unique P-sites corresponding to 3,543 phosphoproteins represented by 24,611 different structures (Table 1). The structures in the database are determined from different methods, including X-ray diffraction (93.85%), NMR (1.48%), EM (<1%), neutron diffraction (<1%), and other combinatorial methods (<1%). Scop3P also contains 78,882 human amino acid variants with disease information obtained from UniProtKB/Swiss-Prot. 80.7% (68478) of these variants are mapped onto 10,416 phosphoproteins. 877 of these variants fall on P-sites, and 245 of these are deleterious variants associated with one or more diseases (Table 1).

### Web Interface and usage of Scop3P

The information in Scop3P can be accessed through the ‘search’ or ‘browse’ options (Fig 1A). The user can search for a protein by UniProt accession number, entry name, protein name, and keywords, or for the results of an entire experimental data set by its ProteomeXchange identifier (ID) ^47^. The results page displays two levels of information: the sequence level, and the structural level (Fig 1A). A quick preview will be displayed in the top panel where all P-sites are mapped in ball and stick notation on the amino acid sequence. The coloring of the ball reflects the frequency of that P-site across the different phosphoproteomics projects from PRIDE. P-sites are colored blue to differentiate the ones that are mapped onto the structure displayed (Fig 1A). A tooltip (triggered when hovering the mouse over the ball) will give additional information such as modification name, number of unique peptides that contain that modification, number of different PRIDE projects in which this P-site is seen, and the mapped position on the PDB structure. A preview on the righthand side of the panel will highlight the P-site on the structure upon hovering (Fig 1A).

Data is also rendered as tables and interactive graphs. The table at the bottom of the sequence annotation gives an overview of the P-sites (Fig 1A), their source (obtained from UniProtKB/Swiss-Prot, from PRIDE, or from both, and, if found in UniprotKB/Swiss-Prot, the evidence information for the P-site (*Experimental*, *By Similarity*, or *Combined* for both). The interactive circular graph contains the residue level predicted secondary structural propensity, backbone dynamics, disorder, and early folding values of the protein in context (Fig 1B). Hovering over a residue in the graph will display the values associated with that residue. As a point of reference, the first amino acid of the protein will be colored dark in all rings of the circular plot.

The phospho-peptide table (Fig 1C) provides information on all PRIDE-derived peptides that contain one or more P-sites for a particular protein. Information such as the ProjectID, peptide sequence, start and end positions of that peptide in the protein sequence, and modified residue position in the protein and peptide sequence are displayed. Hovering over the projectID will display project metadata such as the title, species, submission type, tissues, and publication date. By clicking on the drop-down icon, project information such as project id and frequency of that peptide in the corresponding project can be accessed. In addition, the frequency of P-sites (i.e. the number of times the P-site is seen as phosphorylated or unphosphorylated in particular PRIDE projects) will be shown. This serves as an indication of reliability for that P-site.

In the mutation table, known amino acid variants and their known associated diseases (if any) are given (Fig 1D). These variant details can also be viewed when hovering over the P-site’s ball and stick representation in the top panel.

Finally, in the structure table (Fig 1E), all available structures for the selected protein that contain at least one P-site will be displayed. The idea of visualizing a particular P-site mapped onto multiple protein structures can give insights into the structural context in which the P-site is located in different structural conformations of that protein. In the overview, information such as PDB ID, main chain, interacting chain/molecule, secondary structural propensity, conservation scale, resolution and stoichiometry of the structure, and the position of the P-sites in the PDB structure will be displayed. By clicking on the dropdown icon, secondary structural information such as accessible surface area (ASA), buried surface area (BSA) is given. Upon clicking the PDB ID, a dedicated page for structure will be displayed where the user can color the P-sites based on solvent accessibility, frequency of the site across the different PRIDE projects, and neutral/deleterious polymorphisms (Fig 1E). A complete map of all P-sites mapped on to that particular structure can be viewed upon clicking the “show all P-sites” checkbox.

## Discussion

Scop3P is a user-friendly data resource that allows the analysis of experimentally determined human P-sites in the context of protein structure. It integrates information from different knowledge bases, and shows how re-analysis of large scale public proteomics data sets can add an additional level of significance and confidence to the P-sites based on P-site frequency. Moreover, Scop3P also displays all structural and biophysical information that is available for a particular P-site. This provides additional knowledge about a P-site’s spatial location, structural propensity, and accessibility in the different available conformations of a protein structure. The value of this latter analysis option is highlighted by the fact that 11,399 of the 17,123 P-sites with an available structure have more than one such structure available in Scop3P. Moreover, early folding, disordered propensity, and backbone dynamics data will all provide valuable added information for researchers seeking to understand whether any phospho acceptor residues or related residues in close proximity are crucial for protein folding and stability. Interestingly, by re-processing phosphoproteomics data with the aid of the ionbot search engine (https://ionbot.cloud) we could identify and annotate 1,268 P-sites that are currently annotated as ‘by similarity’ in UniProtKB/Swiss-Prot, thus providing solid experimental evidence for these instances. Moreover, by re-processing we identified and annotated 7,772 proteins that contain at least one P-site which is not annotated as P-site in UniProtKB/Swiss-Prot. Over time, new phosphoprojects in PRIDE will be reprocessed and added to Scop3P, which might add further confirmed or even wholly novel P-sites to the system. The intention is that Scop3P will over time be extended to include other PTMs than can affect residues that can be phosphorylated (e.g., nitrosylation, sulfation, and glycosylation), which might help understand their potential competition for the same residue. Scop3P will be updated and maintained regularly, and will include new data from UniProtKB/Swiss-Prot and PDB, newly reprocessed PRIDE data, and updated data from the various biophysical property predictors. Because of its broad and unique data contents, Scop3P provides a unique and powerful resource to understand the impact of P-sites on human protein structure and function, and can serve as a springboard for researchers seeking to analyze and interpret a given phosphosite or phosphoprotein.

## Data availability

Scop3P is available as a web-interface and can be accessed at https://iomics.ugent.be/scop3p.

## Acknowledgements

PR, LM, WV acknowledge funding from the Research Foundation Flanders under grant agreement number G.0328.16N. LM acknowledges funding from the European Union’s Horizon 2020 Programme under Grant Agreement 823839 (H2020-INFRAIA-2018-1). NH and LM acknowledge funding from a Concerted Research Action grant from Ghent University under grant agreement number BOF12/GOA/014. EV is a postdoctoral research fellow of the Research Foundation Flanders under grant agreement number 12F0816N. The authors are grateful to all submitters to the PRIDE database for making their proteomics data publicly available.

## Author contributions

LM conceived and designed the project. PR performed the data extraction and integration. DT, NT and NH developed the web-interface. WV and EV supervised the work. PR, LM, WV and EV wrote the manuscript.

## Supporting information

Table S1: List of PRIDE projects re-processed

