## Appendix Table S1 for "Scop3P: a comprehensive resource of human phosphosites within their full context"

### Appendix Table S1: PRIDE projects used for re-processing

| | ProteomeXchange ID | Title | Publication date | Submission type | Tissues |
|---|---|---|---|---|---|
| 1 | PXD000474 | Simulated phosphopeptide spectral library for confident site localization | 2015-03-17 | PARTIAL | HeLa cell |
| 2 | PXD000612 | Ultra-deep human phosphoproteome reveals different regulatory nature of Tyr and Ser/Thr-based signaling | 2014-08-06 | PARTIAL | cell culture |
| 3 | PXD000836 | HOPE-fixation of lung tissue allows retrospective proteome and phosphoproteome studies | 2014-05-22 | PARTIAL | lung |
| 4 | PXD001060 | Fe-IMAC column based phospho enrichment | 2015-08-19 | PARTIAL | |
| 5 | PXD001170 | AML_profiling | 2017-02-16 | PARTIAL | blood |
| 6 | PXD001333 | Anion-Exchange Chromatography of Tryptic and Phosphopeptides: WAX vs. SAX and AEX vs. ERLIC | 2015-04-23 | PARTIAL | HeLa cell |
| 7 | PXD001374 | Label-free quantitative phosphoproteomics with novel pairwise abundance normalization reveals synergistic RAS and CIP2A signaling | 2015-08-24 | PARTIAL | HeLa cell |
| 8 | PXD001546 | Reproducibility of label-free single-shot phosphoproteomics applied to CRC cell lines | 2015-04-15 | PARTIAL | cell culture |
| 9 | PXD001550 | Human CRC cell line baseline phosphoproteomics | 2015-04-15 | PARTIAL | cell culture |
| 10 | PXD000089 | Proteomic and phosphoproteomic data of colorectal cancer tissues and cells for Chromosome-Centric Human Proteome Project | 2013-01-11 | PARTIAL | - |
| 11 | PXD001565 | Evaluation of phospho-tyrosine antibodies for label-free phosphoproteomics | 2015-12-09 | PARTIAL | colon,brain |
| 12 | PXD002255 | Enrichment strategy for searching missing protein | 2015-07-14 | COMPLETE | cell culture,blood serum |
| 13 | PXD002286 | Phospo-proteomic profiling of Castration Resistant Prostate Cancer | 2016-08-19 | PARTIAL | |
| 14 | PXD002394 | Proteomic and phosphoproteomic analysis of cisplatin resistance in patient derived serous ovarian cancer | 2017-05-02 | PARTIAL | cell suspension culture |
| 15 | PXD002646 | A systematic evaluation of nanocast metal oxide microspheres for phosphoproteomics applications | 2017-05-17 | PARTIAL | |
| 16 | PXD002990 | Complementary Phosphoproteomic Approaches Link IL-23R Downstream Signaling with Metabolic Adaptation in Lymphocytes | 2016-04-21 | COMPLETE | lymph node |
| 17 | PXD003198 | Characterisation of pancreatic ductal adenocarcinoma subtypes by global phosphotyrosine profiling | 2016-06-09 | PARTIAL | pancreatic cell line |
| 18 | PXD003531 | Proteomics of Primary cells derived from Ovarian Cancer | 2017-04-03 | PARTIAL | primary cell |
| 19 | PXD004252 | Modulating the selectivity of affinity absorbents to multi-phosphopeptides by a novel competitive substitution strategy | 2016-08-03 | PARTIAL | cell culture |
| 20 | PXD004415 | Quantitative phosphoproteome analysis of cisplatin-induced apoptosis in Jurkat T cells | 2017-05-11 | COMPLETE | - |

| | | | | | |
|---|---|---|---|---|---|
| 21 | PXD004447 | Mirroring the charged termini; ETD/HCD fragmentation characteristics of LysargiNase and tryptic peptides and their benefits for peptide sequencing in proteomics | 2016-12-02 | PARTIAL | cell culture |
| 22 | PXD004452 | HeLa proteome of 12,250 protein-coding genes | 2017-06-12 | PARTIAL | liver,colon |
| 23 | PXD004940 | Performance of the Orbitrap Fusion Lumos Tribrid in single-shot analyses of human samples | 2017-03-09 | PARTIAL | HeLa cell |
| 24 | PXD005366 | Robust, sensitive and automated phosphopeptide enrichment optimized for low sample amounts applied to primary hippocampal neurons | 2016-12-14 | PARTIAL | - |
| 25 | PXD006482 | Identification of Missing Proteins in the Phosphoproteome of Kidney Cancer | 2017-09-01 | COMPLETE | kidney |
| 26 | PXD006114 | Phosphoproteomics of in vivo resistance to EGFR-targeted therapy in lung cancer cells | 2017-08-28 | COMPLETE | cell culture |
| 27 | PXD003215 | A Novel Method for Isolating Whole Protein from Human Cranial Bone | 2016-10-05 | COMPLETE | cranium |
| 28 | PXD003657 | Antibody-independent identification of bovine milk-derived peptides in breast-milk | 2016-07-19 | COMPLETE | |
| 29 | PXD003660 | Early Signaling Dynamics of EGFR | 2016-06-24 | COMPLETE | cell culture |
| 30 | PXD003709 | Preferential phosphorylation on old histones during early mitosis in human cells | 2016-06-01 | COMPLETE | cell culture |
| 31 | PXD002057 | Proteomic analysis identifies novel pathways linking epithelial-to-mesenchymal transition with resistance to HER2-targeted therapy | 2016-02-22 | COMPLETE | cell culture |
| 32 | PXD002436 | Monitoring cellular phosphorylation signaling pathways into chromatin and down to the gene level | 2015-11-09 | COMPLETE | HeLa cell |
| 33 | PXD000964 | MS approach assessment and Mad1 phosphopeptide analysis | 2014-10-31 | COMPLETE | cell culture |
| 34 | PXD000680 | Stable isotope labeling of phosphoproteins for large-scale phosphorylation rate determination | 2014-04-15 | COMPLETE | HeLa cell,HEK-293 cell |
| 35 | PXD000218 | System-level analysis of cancer and stomal cell specific proteomes reveals extensive reprogramming of phosphorylation networks by tumor microenvironment | 2014-04-01 | COMPLETE | colon |
| 36 | PXD000674 | PeptideShaker | 2014-01-27 | COMPLETE | HeLa cell |
| 37 | PXD000021 | D-score | 2013-01-23 | COMPLETE | - |
